# Simultaneous Determination of Protein Structure and Dynamics Using Cryo-Electron Microscopy

**DOI:** 10.1101/219972

**Authors:** R. Pellarin, M. Vendruscolo

## Abstract

Cryo-electron microscopy is rapidly emerging as a powerful technique to determine the structures of complex macromolecular systems elusive to other techniques. Since many of these systems are highly dynamical, characterising also their movements is a crucial step to unravel their biological functions. To achieve this goal, we report an integrative modelling approach to simultaneously determine structure and dynamics from cryo-electron microscopy density maps. By quantifying the level of noise in the data and dealing with their ensemble-averaged nature, this approach enables the integration of multiple sources of information to model ensembles of structures and infer their populations. We illustrate the method by characterising structure and dynamics of the integral membrane receptor STRA6, thus providing insights into the mechanisms by which it interacts with retinol binding protein and translocates retinol across the membrane.

## Introduction

Cryo-electron microscopy (cryo-EM) (1-9) is a powerful structural biology technique that enables the characterization of complex biological systems that are not readily amenable to other traditional techniques, such as X-ray crystallography and nuclear magnetic resonance spectroscopy. With the continuous progress in the development of both instrumentation and image processing software, cryo-EM is rapidly approaching the resolution of X-ray crystallography (2-9), allowing the structures of complexes or individual proteins of great biological relevance to be determined in their natural environments at nearly-atomistic resolution (10-17).

Despite these great advances, a major challenge remains open – the characterization of the structural dynamics of the systems observed (18, 19). This is a crucial problem, since most macromolecular machines populate multiple conformational states, and their functions depend on the interplay between structure and dynamics. In several cases, cryo-EM can detect alternative conformations (3-6, 15, 16), provided that two-dimensional (2D) images of particles with distinct conformations are separately classified (5, 6, 20). However, if the system has highly dynamic regions or the low resolution of 2D images hinders classification, three-dimensional (3D) reconstructions generated by combining multiple class-averages present regions at lower resolution. These regions may correspond to flexible parts of the system, which are averaged out in the class-averaging and reconstruction process, to particularly noisy portions of the map caused for example by radiation damage, or to both.

Several modelling approaches are currently used to translate cryo-EM data into structural models (21, 22). Following the classification proposed in Ref. (22), these approaches include methods for rigid-body fitting (Chimera (23), COAN (24), EMfit (25), Modeller (26), MultiFit (27), SITUS (28)), flexible fitting (EMFF (29), MDFF (30), Modeller (26), SITUS (28), MDFIT (31), Flex-EM (32)), homology modelling (Fold-EM (33), ROSETTA (34), Modeller (26)), *de novo* modelling (EM-fold (35), SITUS (28), IMP (36), RELION (20), Phenix (37)), and integrative approaches (IMP (36)). All these approaches search for single structural models that minimize the deviation between observed and predicted cryo-EM density maps, usually by incorporating additional restraints into Monte Carlo or molecular dynamics (MD) simulations. Some of these technique can provide multiple alternative models that individually fit the input map to a certain extent (24, 38), in the same way as multiple molecular models are derived from nuclear magnetic resonance (NMR) spectroscopy data (39), and routinely deposited in the PDB database. The sets of these models can be considered as ‘uncertainty ensembles’, since they reflect the limited information available on the systems and thus the fact that different models might be equally consistent with the input data; they do not, however, reflect the conformational heterogeneity arising from the internal dynamics of the systems themselves (40-44). In many cases, as for instance in inferential structure determination (45), weights can be associated to the members of these uncertainty ensembles. These weights quantify the consistency with the input information and thus the overall confidence in each individual model (46); they are not, however, statistical weights, since they are not meant to represent the equilibrium populations of different conformations present in a conformationally heterogeneous mixture.

The methods described above can successfully determine single-structure models as well as uncertainty ensembles from cryo-EM data, but they do not to provide information about the thermodynamics and dynamics of the systems studied. Such information can only be extracted from ‘thermodynamic ensembles’, which represent sets of conformations, together with their statistical weights, that coexist in a heterogeneous mixture along with their equilibrium populations (40, 44, 47). To obtain a thermodynamic ensemble, one should search for structural models whose ensemble-averaged observables, rather than individual conformations, fit the input data (40, 44, 47). Importantly, in the construction of such thermodynamic ensembles, all possible sources of errors should be taken into account when evaluating the fit of the ensemble with the experimental data (48).

Here, we report an approach to enable the simultaneous determination of structure and dynamics of proteins and protein complexes by modelling thermodynamic ensembles from cryo-EM density maps. This approach is based on the recently introduced metainference (46), a general Bayesian inference method (45) that enables the modelling of heterogeneous systems from noisy, ensemble-averaged experimental data. This method incorporates a Bayesian treatment of cryo-EM data that accounts for variable level of noise in the maps and enables structural modelling at atomistic resolution (49).

## Materials and Methods

### Overview of the metainference approach

Metainference (46) is a Bayesian framework (45) for modelling thermodynamic ensembles by integrating prior information on a system (physical, chemical or statistical knowledge) with noisy experimental data. The method is designed to deal with conformationally heterogeneous systems, in which experimental observations reflect an ensemble of states rather than a single conformation, and with data affected by known and unknown errors. The metainference approach enables: *i)* optimally combining and weighting multiple sources of information; *ii)* modelling thermodynamic ensembles; *iii)* determining the population of all states in the ensemble; *iv)* determining the level of noise in the input data. In doing so, metainference allows overcoming the limitations of individual computational and experimental technique (40). Standard MD simulations, which are hampered by inaccuracies in the empirical physico-chemical description of the system (the force field), will be complemented by experimental data, which alone would provide only sparse information affected by random and systematic errors.

In metainference, the generation of models is guided by the metainference energy function, defined as *E*_*MI*_ = −*k*_*B*_*T* log*p_MI_*, where *k*_*B*_ is the Boltzmann constant, *T* the temperature of the system, and *p*_*MI*_ the metainference posterior probability. The posterior expresses the probability of observing a given set of structural models, and possibly other parameters, in terms of prior information and data likelihood. The former encodes physico-chemical knowledge of the system; the latter quantifies the agreement with experimental data and incorporates a model of experimental noise. In case of Gaussian noise, the metainference posterior of observing a set of *N* conformations ***X*** = [*X*_*r*_] given *N*_*d*_ independent experimental data points ***D*** = [*d*_*i*_] is:

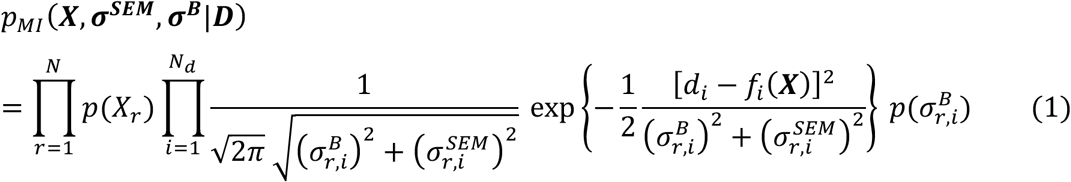

where 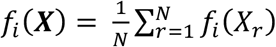 is the average of the predictor (or *forward model*) *f*_*i*_ for the experimental observable *d*_*i*_ calculated over the set of *N* conformations, 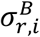 a parameter quantifying the noise level in data point *d*_*i*_, and 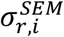 the statistical error in calculating an ensemble average of *f*_*i*_ using a finite set of conformations. 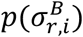 and *p*(*X*_*r*_) are the priors on 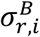 and *X*_*r*_, respectively.

A sample of the posterior distribution is typically obtained by running a multiple-replica simulation (50) guided by the associated metainference energy function:

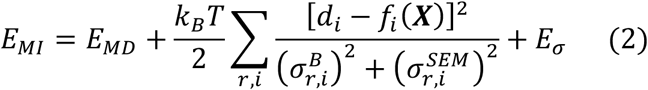

in which the force field of standard MD simulations, 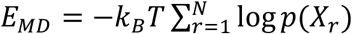, is modified by *i)* a series of (harmonic) *data restraints* that ensure the agreement of the structural ensemble with the experimental data and *ii)* a series of *error restraints*, 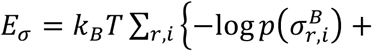 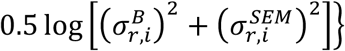. In this multi-replica simulation scheme, one needs to sample, for each replica, not only the space of possible conformations *X*_*r*_, but also of the parameters 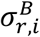 that quantify the level of noise in the data. This is typically achieved by a Gibbs sampling scheme, which combines MD to sample the coordinates space with Monte Carlo for the noise parameters 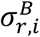. The values of these noise parameters ultimately determine the intensities of the harmonic data restraints: low noise will result in a strong structural restraint, inconsistent data points and outliers will be automatically labelled as *noisy* and down-weighted in the construction of the final ensemble.

### The metainference approach for cryo-EM density maps

In the following, we define the metainference components, previously introduced in general terms, specifically for the case of cryo-EM data. The development of these elements builds on the approach proposed in Ref. (49). A summary of all the elements at the basis of our approach is reported in **Table S1**. The method is implemented in the PLUMED-ISDB module (51) of the open-source, freely-available PLUMED library (www.plumed.org) (52).

#### Experimental data

Typically, a cryo-EM density is distributed as a map defined on a grid, or set of voxels, in real space. In our approach, we will represent the experimental density map using a Gaussian Mixture Model (GMM) *ϕ_D_* with *N*_*D*_ components (data-GMM):

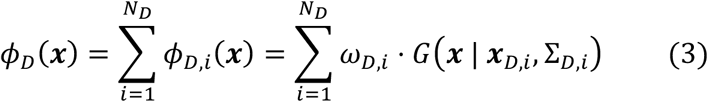

where *ω*_*D,i*_ is the (normalized) weight of the *i*-th component of the data GMM and *G* a normalized Gaussian function centered in ***x***_*D,i*_ and with covariance matrix Σ_*D,i*_. This representation has several advantages. First, it is computationally convenient to use an analytical representation of the input data, rather than a discrete definition on a grid. Second, a GMM can provide a representation of the data in terms of independent bits of information, while in the grid representation neighboring voxels should be considered as correlated data points affected by correlated noise. Finally, a GMM can be tuned to represent the data at different resolutions, from coarse-grained for initial efficient modelling or for low-resolution maps, to atomistic for refinement of high-resolution maps. In order to efficiently fit high-resolution maps at near-atomistic detail, we used a divide-and-conquer approach (49), which starts from a low-resolution fit using few Gaussians and refines it in subsequent iterations to increase the resolution of the final GMM (Supplementary Information).

#### The forward model

We developed a forward model to simulate a cryo-EM map from a structural model. As for the representation of the experimental map, the forward model *ϕ_M_* is a GMM (model-GMM). Since here we employed high-resolution synthetic and real cryo-EM maps, we represented each heavy atom of the system by one Gaussian function, whose parameters were obtained by fitting the electron atomic scattering factors (53) for a given atomic specie (Supplementary Information, **Table S2**). In case of low-resolution maps, a single Gaussian can be used to represent each coarse-grained bead, with the Gaussian width proportional to the size of the bead (49).

#### The noise model

The deviation between predicted and observed density maps is measured in terms of the overlap *ov*_*MD,i*_ between the model-GMM *ϕ_M_* and the *i*-th component *ϕ*_*D,i*_ of the data-GMM

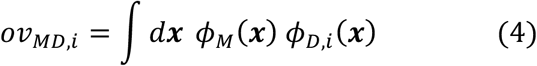

The overlap *ov*_*MD,i*_ can be expressed in a computationally-convenient analytical form (49), being *ϕ*_*D,i*_(***x***) a Gaussian function and *ϕ*_*M*_(***x***) a GMM. In a heterogenous system, the forward model *ϕ*_*M*_ is an average over the *N* metainference replicas 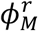, and thus the overlap can be written as

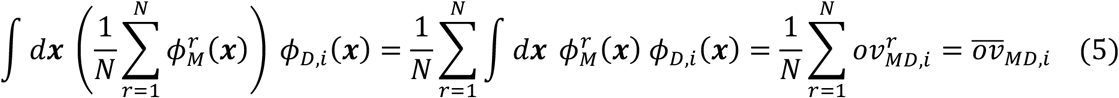

For each component of the data-GMM, we used a Gaussian noise model with one uncertainty parameter per data point to account for variable level of noise across the map. The data likelihood for the overlap *ov*_*DD,i*_ of the *i*-th component of the data-GMM with the entire data-GMM can then be written as:

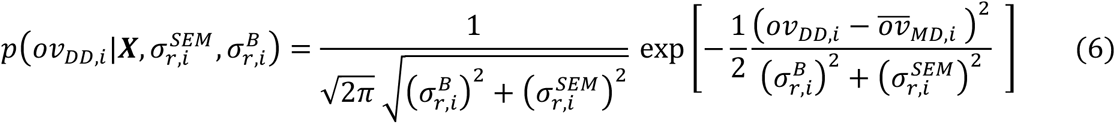

#### Prior information

As structural prior information *p*(*X_r_*), we used the AMBER99SB*-ILDN molecular mechanics force field (54) and the GBSA implicit model of water (55). For the uncertainty parameters, we used an uninformative Jeffreys prior (56). To avoid sampling all the uncertainty parameters, we marginalized them prior to simulating the system (Supplementary Information). The level of noise in each component of the data GMM can then be estimated *a posteriori* using all the structural models produced by the metainference simulations (Supplementary Information).

#### Metainference energy function

After defining the noise model as outlined in the previous paragraphs and marginalizing all the noise parameters, the metainference energy function for cryo-EM data becomes:

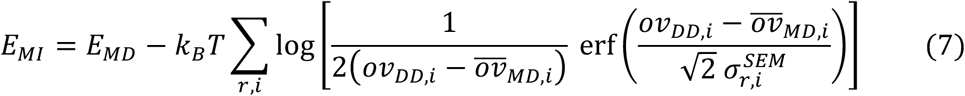

where erf(*x*) is the error function.

### General details of the metainference simulations

All simulations were performed using as prior the AMBER99SB*-ILDN force field (54) along with the GBSA implicit model of water (55). This is a computationally convenient combination of force fields, although its accuracy has been shown to be modestly inferior compared to AMBER99SB*-ILDN combined with explicit solvent models (57). Starting models were equilibrated at 300K for 1 ns. A time step of 2 fs was used together with LINCS constraints (58). Van der Waals and Coulomb interactions were cut off at 2.0 nm. Neighbour lists for all long-range interactions were cut off at 2.0 nm and updated every 10 MD steps. Simulations were carried out using a non-periodic cell in the canonical ensemble at 300K, enforced by the Bussi-Donadio-Parrinello thermostat (59). Configurations were saved every 2 ps for post-processing. To improve computational efficiency, the cryo-EM restraint was calculated every 2 MD steps (60), using neighbor lists to compute the overlaps between model and data GMMs, with cutoff equal to 0.01 and update frequency of 100 steps. Well-tempered metadynamics (61) was used to accelerate sampling of the metainference ensemble (Supplementary Information). All simulations were carried out with GROMACS 4.5.7 (62) and PLUMED (52). Parameters of the GBSA implicit solvent were imported from GROMACS 5.1.4.

### GroEL metainference simulations

The crystal structures of apo GroEL (PDB code 1XCK) (63) and GroEL-ADP complex in the relaxed allosteric state (PDB code 4KI8) (64) were used to generate a synthetic cryo-EM map, using the following procedure. Chains A were extracted from the two pdbs (**Figure 1A**) and aligned using UCSF Chimera (23). MODELLER v9.17 (26) was used to generate a comparative model (GroEL-ADP*) of the sequence of apo GroEL based on the GroEL-ADP complex template (64). The gmconvert utility (65) was then used to separately convert the apo-GroEL and GroEL-ADP* atomistic models into two density maps (**Figure 1B**). Radius and weight for the conversion were determined by the “residue type” method implemented in gmconvert. The final synthetic map was computed as the average of the two individual maps, with equal weight (**Figure 1C**). A divide-and-conquer approach (49) was used to fit a GMM with 4000 components, which resulted in a cross-correlation with the original map of over 0.99 (**Figure S1**). Initial models for the metainference production run were randomly extracted from the 1 ns-long equilibration run initiated from the apo-GroEL model. The metainference ensemble was simulated using 4 replicas for a total aggregated time of 50 ns. 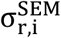 was kept constant for all replicas and set to 0.01 ov_DD,i_. This parameter determines the maximum intensity of the cryo-EM restraint in case of absence of data noise 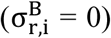 and was set to the minimum value that allowed a proper integration of the cryo-EM restraint. To enhance sampling, we used well-tempered metadynamics with W_0_ = 1000 kJoule/mol and γ = 150,000.

**Figure 1.**
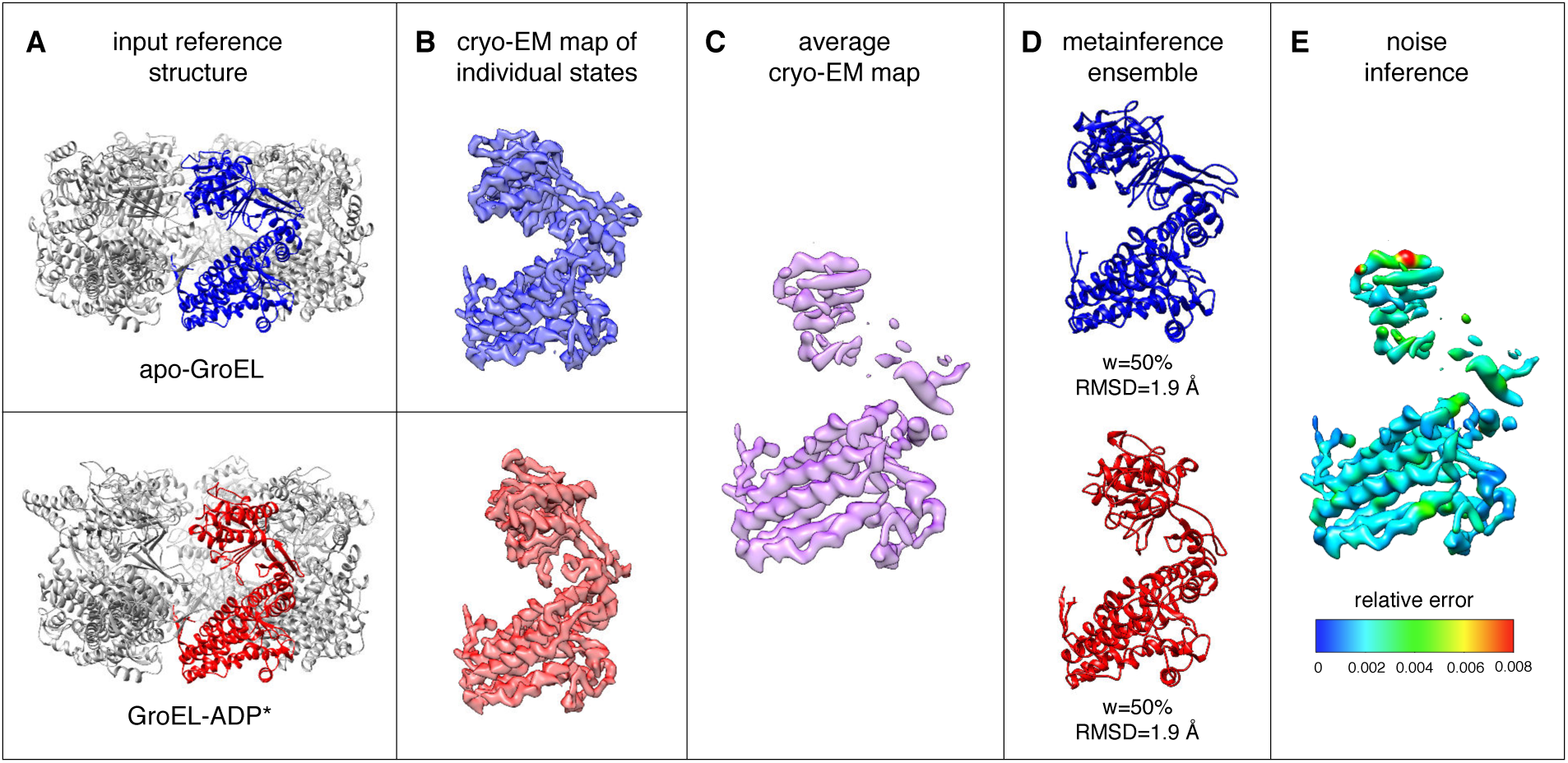
Validation of the metainference approach on a conformationally heterogeneous ensemble of the chaperonin GroEL. The crystal structure of Apo-GroEL (63) (A, blue) and a comparative model built from the structure of GroEL in complex with ADP (64) (A, red) were used to create synthetic cryo-EM maps at near-atomistic resolution (B). An average map was then computed by mixing contributions from the two models in ratio 1:1 (C). The metainference approach was capable of disentangling the contribution of the two states (D), determining their relative populations in the mixture, and inferring the local level of noise in the map (E).

### STRA6 metainference simulations

The cryo-EM map of the complex of zebrafish STRA6 with co-purified calmodulin at 3.9 Å resolution (EMD code 8315 (66)) was fit with a GMM using a divide-and-conquer approach (49). The final GMM was composed of 11585 Gaussians and resulted in a cross-correlation with the original map equal to 0.97 (**Figure S2**). The cytosolic loop (residues 575-597) missing from the deposited model (PDB code 5SY1) was modelled using MODELLER v9.17. The residue numbering scheme was kept as in the deposited model. The resulting comparative model was then equilibrated at 300 K, as previously described. Initial models for two independent production runs (labelled as RUN I and RUN II) were then randomly extracted from the equilibration run. The metainference ensemble was simulated using 16 replicas for a total aggregated time of 355 ns per production run. 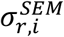 was set to 0.1. *ov*_*DD,i*_. To enhance sampling, we used well-tempered metadynamics with *W*_0_ = 5000 kJoule/mol and *γ* = 950,000. Details of the analysis of the simulations (stereochemistry assessment, comparison with the experimental cryo-EM map, free-energy calculations, noise inference, and convergence analysis) are reported in the next sections.

### Stereochemistry assessment of the STRA6 ensemble

To measure the stereochemical quality of the ensemble of STRA6 models generated by the metainference method, we calculated the distribution of the backbone dihedral angles *ϕ* and *ψ* on the conformations sampled in the two independent simulations. To achieve this, we used PROCHECK (67), specifically the *procheck_nmr* collection of codes designed to evaluate the quality of NMR ensembles. This program classifies all residues in all models in 4 regions of the Ramachandran plot (**Figure S3A**): residues in most favoured regions (red), in additional allowed regions (yellow), in generously allowed regions (light yellow), and in disallowed regions (white). The percentages of residues in each of these regions for the two independent metainference simulations were, in both cases: 87.4%, 11.6%, 0.5%, and 0.5% (**Figure S3C**). These values were comparable to those obtained using the STRA6 deposited model (PDB code 5SY1): 86.4%, 13.3%, 0.3%, 0.0% (**Figure S3B**).

### Comparison with the STRA6 experimental cryo-EM map

To evaluate the quality of the fit of our metainference ensemble with the experimental map and compare it with the deposited model, we used the *gmconvert* utility (65) to calculate synthetic cryo-EM maps from structural models. It is important to note that the algorithm implemented in *gmconvert* is different from our forward model and from the approach implemented in RELION (20) and used in Ref. (66) to refine the deposited model. In this way, *gmconvert* provides a method to predict a cryo-EM map independent from those used in the generation of our ensemble and in the refinement of the deposited model. We believe that this is a fair procedure to evaluate the agreement with the experimental map. The local cross-correlation (CC) with the experimental map (EMD code 8315) of the average map computed on our metainference ensemble and the one computed on the deposited model was evaluated in a 5 voxel sliding window using the “*vop localcorrelation*” command in UCSF Chimera (23). The same program was then used to color the metainference maps and deposited-model maps based on the value of the local CC (**Figures 2B,D** and **S4B,D**).

**Figure 2.**
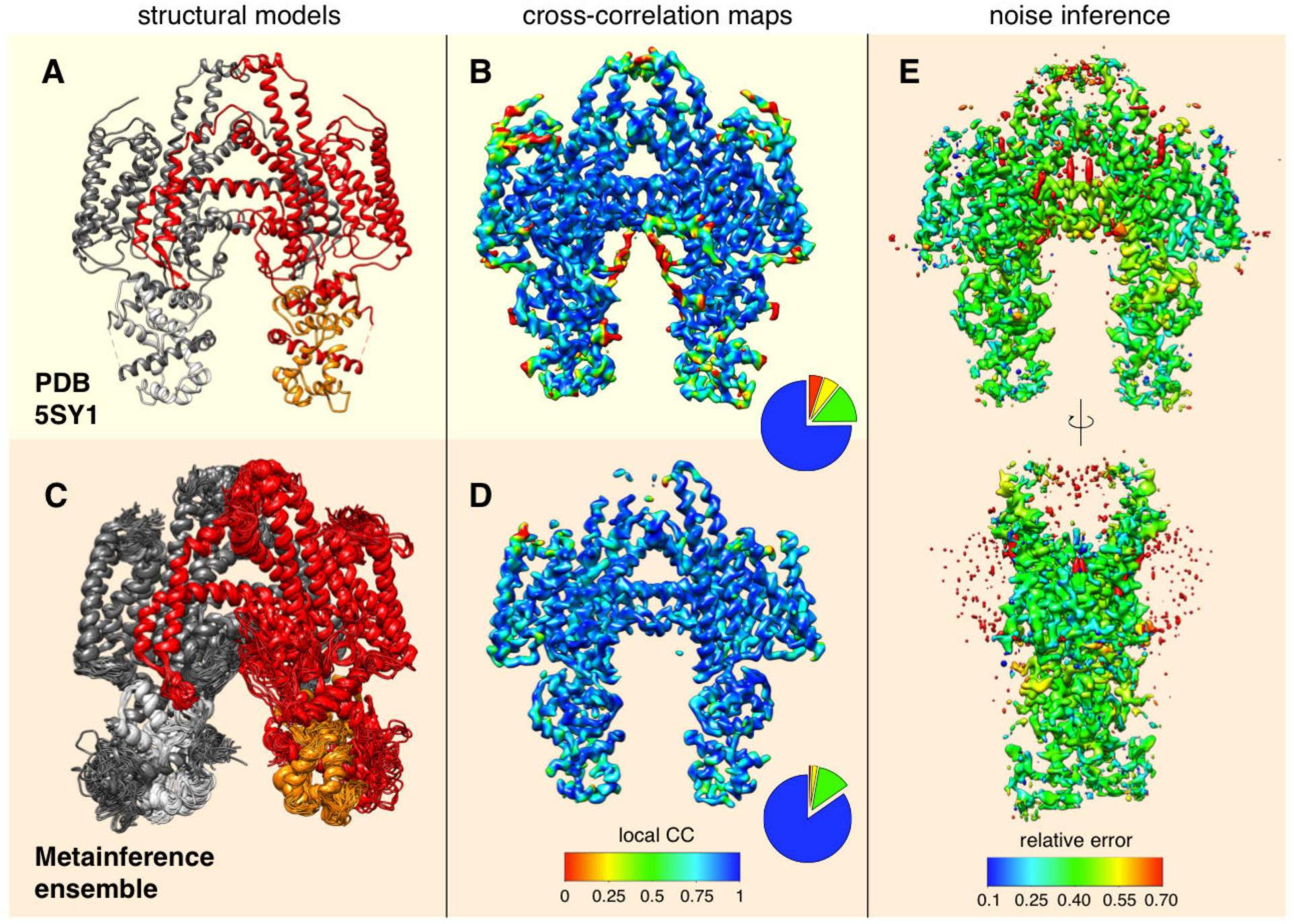
Structure, dynamics, and noise characterization of the STRA6 membrane complex. Compared to the single-structure model (PDB 5SY1, A), the metainference ensemble displays a higher degree of flexibility (C). We calculated the predicted cryo-EM maps from the single-structure model (B) and metainference ensemble (D) and evaluated the global and local cross-correlation (CC) with the experimental map. The metainference map provides a better CC with the experimental map (global CC=0.91) compared to the single-structure map (global CC=0.86), especially in the more dynamical regions of STRA6. Cross-correlation maps are visualized at a threshold of 3.5. The pie charts report the distributions of local CC in the regions of the single-structure and metainference maps with density between 3.4 and 3.6. The level of relative error in the experimental map inferred by metainference is rather uniform, with the exception of the regions occupied by cholesterol and amphiphols (E).

### STRA6 thermodynamic ensemble analysis

To characterize the thermodynamics of the metainference ensemble obtained from the STRA6 cryo-EM data, we projected all the conformations onto a set of structural descriptors, or collective variables (CVs). To shed light into the dynamics of the external cleft and the binding of RBP, the CVs were defined as: a) the distance between the Cα atoms of residues L323 and N441 in the first chain of STRA6 and b) the distance between the Cα atoms of the corresponding residues in the other identical chain (L323’ and N441’). To investigate the role of the JM helix in retinol binding and release, the CVs were defined as: a) the distance between the geometric centers of residues P248-D252 in JM and V535’-F538’ in JML and b) the distance between the geometric centers of residues P248-D252 in JM and L366-R376 in TM7. The PLUMED *driver* utility (52) was used to calculate the CVs defined above from the metainference ensemble, which where then used to construct the associated free energies (**Figures 3B** and **4C**).

**Figure 3.**
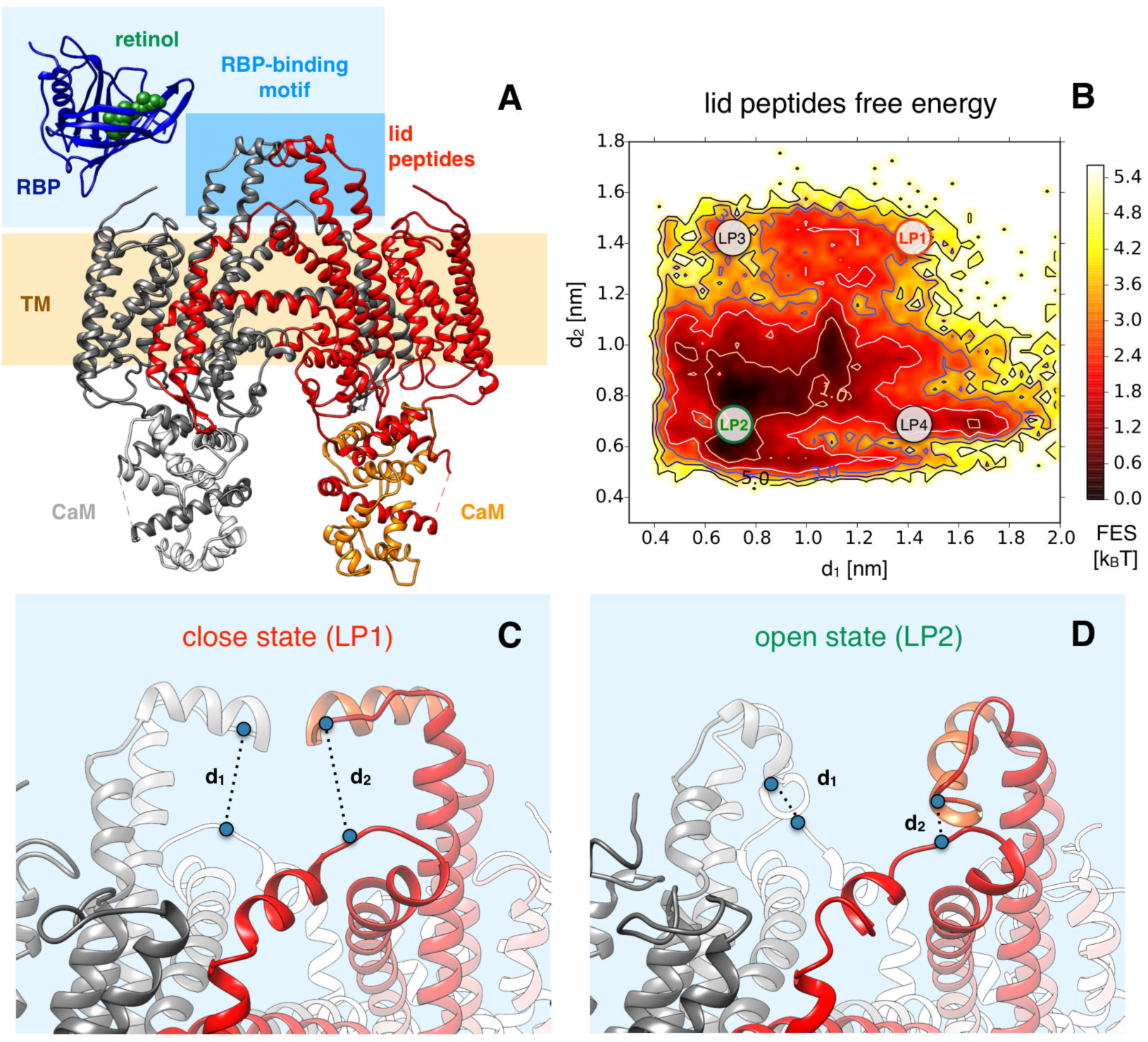
Structural insights into the mechanism of RBP binding. To obtain insight into the mechanism of RBP binding (A), we projected all conformations of the metainference ensemble along the two collective variables d1 and d2, which were defined as the distances between residues N441 in the TM8-TM9 loop and L323 in LP, in each of the two identical monomers. The resulting free energy landscape indicates an equilibrium among different conformations (B). The ‘close’ state observed in the single-structure model (LP1, C), in which the two LPs are close together, has a relatively low population. A more stable state is an ‘open’ conformation in which the two LPs fold back to interact with the TM8-TM9 loop (LP2, D). States in which only one of the two LPs folds back are also visible (LP3 and LP4, B).

**Figure 4.**
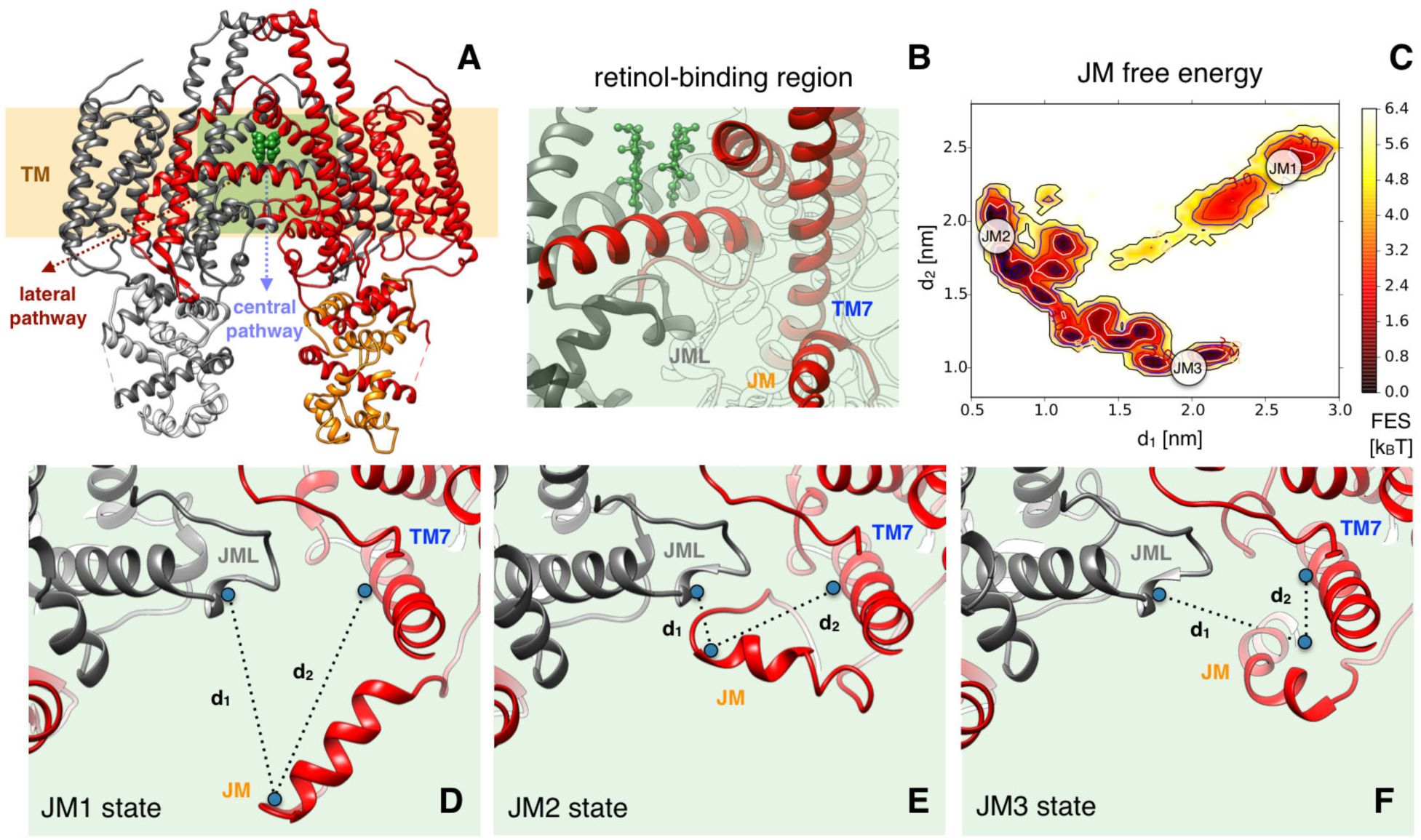
Structural insights into the mechanism of retinol release. To investigate the role of JM in retinol binding and release (A,B), two collective variables were defined as the distance between the geometric centers of residues P248-D252 in JM and V535’-F538’ in JML (d1) and the distance between the geometric centers of residues P248-D252 in JM and L366-R376 in TM7 (d2). The associated free energy landscape indicated an equilibrium among different conformations (C). JM, which in the PDB model resides far apart from JML and TM7 (JM1, D), can transiently iteract with both JML (JM2, E) and TM7 (JM3, F), suggesting a possible role of JM in facilitating retinol release by weakening the JML-IM interaction and the stability of the binding site situated between the IM helices (B).

### Convergence of the STRA6 metainference simulations

To assess convergence, we performed a cluster analysis of the two independent STRA6 simulations. We first merged the two runs (labelled as RUN I and RUN II) and then performed a cluster analysis on the concatenated trajectory, after discarding the initial 20% of each run. We used the GROMOS clustering algorithm (68) and the backbone RMSD as measure of conformational similarity, with a cutoff equal to 0.35 nm. In this way, we defined a discrete set of 18 conformational states of STRA6 common to the two production simulations. To assess the convergence of a given run, for each cluster we calculated its population and error as the average and standard deviation of the population computed in the first and second half of each simulation, and then we compared the results from the two independent runs (**Figure S10**). Additional characterization of the STRA6 ensemble obtained from RUN I and RUN II is reported in **Figures S4**, **S7** and **S8**.

## Results and Discussion

### Validation with GroEL data

We first assessed the accuracy of the metainference approach by applying it to the case of the chaperonin GroEL in two different conformations (**Figure 1A**), apo-GroEL, the compact crystal structure of apo GroEL (63), and GroEL-ADP*, a comparative model built from a an extended allosteric state adopted by GroEL in complex with ADP (64). We created synthetic cryo-EM maps from these two models (**Figure 1B**), and used them to construct an average cryo-EM map that equally mixed contributions from the two states (**Figures 1C**), which have a backbone root-mean-square deviation (RMSD) of 3.9 Å (**Figure S5**). As expected, the metainference ensemble was characterized by the presence of two distinct, equally populated structural clusters (**Figures 1D**), which corresponded to apo-GroEL and GroEL-ADP*, with backbone RMSD of the centers of the two clusters from the correspondent model equal to 2.0 Å (**Figure S5**). Our approach accounts for unkown and variable level of noise across the experimental map during the modelling of the ensemble. In this case, the only source of error, i.e. of deviation between predicted and observed maps, is the difference between the procedures used to generate the synthetic map and in our modelling approach. The inferred level of relative noise was fairly uniform across the map, with average value of about 0.003 (**Figures 1E** and **S6**).

### Structure and dynamics of the STRA6 membrane receptor

We then applied our metainference approach to the integral membrane receptor STRA6 (66), which mediates the cellular uptake of retinol by extracting it from its carrier (retinol binding protein, or RBP) and moving it across the membrane (69). Recently, zebrafish STRA6 was determined at 3.9 Å resolution by single-particle cryo-EM (EMD code 8315) (66). The structural model obtained from the cryo-EM map (PDB code 5SY1) revealed a dimer of STRA6 in complex with the protein calmodulin, and enabled the characterization of the regions involved in RBP binding and retinol translocation across the membrane (66). In the present work we focus in particular on the regions of the STRA6 map that were determined at lower resolution (66), including the periphery of the complex and the RBP binding region, in order to show that they can be interpreted in terms of a combination of conformational dynamics and noise in the data.

#### The STRA6 metainference ensemble

Starting from the 5SY1 structure (**Figure 2A**), we modelled a thermodynamic ensemble of conformations by integrating cryo-EM data (**Figure S2**) with an *a priori,* physico-chemical knowledge of the system (Materials and Methods). In the generated ensemble (**Figures 2C** and **S4C**), STRA6 presents a significant degree of flexibility around the single-structure model, in particular in the N-terminal and C-terminal domains, the lid peptide (LP) and the RBP-binding motif, the cytosolic loop, and the juxtamembrane (JM) helix (**Figure S7A**). These regions correspond to areas at lower resolution in the experimental map (66), as well as in the back-calculated maps (**Figure 2B,D** and **S4B,D**). The cytosolic loop, which is not included in the 5SY1 structure, displays the largest fluctuations (**Figure S7A**, magenta), while calmodulin (**Figure S7B**) and the STRA6 trans-membrane (TM) domain (**Figure S7A**, grey) are instead more rigid and deviate less from the 5SY1 model (**Figure S7C,D**). The latter result is particularly relevant, given the low accuracy of the prior in the TM region (Materials and Methods).

We then measured the agreement of the metainference ensemble with the experimental data by calculating the maps predicted from the ensemble and the single-structure model (Supplementary Information). We found that the metainference ensemble provided a better cross-correlation with the experimental map (global CC=0.91) than the single-structure model (global CC=0.86), especially in the more dynamical regions of STRA6 (**Figures 2B,D** and **S4B,D**). We also verified that the improved agreement with the experimental data was not achieved at the expenses of the stereochemical quality of the models (**Figure S3**).

#### Noise inference

To quantify the level of noise in the data, we calculated an error density map from the metainference ensemble (Supplementary Information) and visualized it onto the experimental map (**Figures 2E and S4E**). The inferred level of relative error was fairly uniform, with average value of about 0.38 (**Figure S8**), except for few specific regions: the binding sites in-between the two horizontal intramembrane helixes (IM), the interior of the outer cleft, and the external region surrounding the TM domain (**Figures 2E and S4E**). It has been suggested (66) that all these regions are occupied by components that were not explicitly modeled, including cholesterol in the binding sites and in the outer cleft, and amphiphols in the shell around the TM domain. Consistently with the conclusion that these electron densities could not be explanined by the presence of STRA6 alone, the metainference method resulted in the assignement of a high level of noise to these regions.

#### Dynamics in the RBP-binding region

Next, we investigated how the conformational dynamics obtained by the metainference approach can be linked to the biological function of STRA6 and thus to which extent the resulting ensemble can be more informative and predictive than a single-structure model. The observed flexibility in the N-terminal domain might be functional, as this region might act as a sensor for unidentified ligands (66) or to facilitate RBP recognition and recluting through non-specific interactions, prior to binding specifically. We found that the LP region (**Figure 3A**) is characterized by the presence of an equilibrium among different conformations (**Figure 3B**). The state in which the two LPs from the two STRA6 monomers are close together (LP1, **Figure 3C**), as in the PDB model, appear to have a relatively low population. In the most populated state, the two LPs fold back and approach the loop region between TM8 and TM9 (LP2, **Figure 3D**). States in which only one of the two LPs folds back were also present (LP3 and LP4, **Figure 3B**). As these results are consistent with the possibility that LP2, rather than LP1, may be more productive for RBP binding, incorporating further information about the physico-chemical environment in the surroundings of the LP region will improve their prior description and offer a more accurate quantification of their relative stabilities. The existence of the LP2 state, not directly visible in the single-structure model, could explain the fact that inserting a Myc tag at the apex of the TM8-TM9 loop impairs RBP binding (70). This effect would be the result of either altering the equilibrium among LP states in favor of a conformation not productive for binding or destabilizing the actual binding state, depending whether LP2 is the actual binding conformation.

#### Translocation of retinol across the membrane

Concerning the exit mechanism of retinol from STRA6 into the cytosol (**Figure 4A,B**), the single-structure model suggests a lateral pathway, as the alternative pathway from the central dimer interface would require significant conformational changes (66). From our study of the conformational dynamics of STRA6, we identify a potential role of the JM helix in regulating retinol release through either of the two pathways. This fragment populates multiple distinct confomations in our ensemble (**Figure 4C**). In one state (JM1, **Figure 4D**), JM points outwards from STRA6, as in the single-structure model. In the second (JM2, **Figure 4E**), this peptide interacts with the JM loop (JML), which is situated below the horizontal IM helices and the putative retinol binding site (**Figure 4B**). In another state (JM3, **Figure 4F**), JM resides in proximity to TM7. This complex equilibrium among states suggests that JM can transiently interact with JML and possibly compete with the formation of the conserved salt bridge D539-R511’ between JML and IM. This salt bridge appears to be crucial for consolidating the JML-IM interaction and stabilizing the retinol binding site located in-between the IM helices, as its disruption by mutation in human STRA6 results in Matthew-Wood syndrome (71). By competing with the salt bridge formation, JM could promote IM and JML mobility, weaken the stability of the retinol binding site, and eventually favour retinol unbinding. Translocation of retinol across the membrane can later occur through either the lateral or central pathways. The latter scenario would require additional conformational changes, which might be facilitated by the transient disruption of the salt bridge and the increased mobility of this region.

This dynamical picture of JM offers interesting perspectives, regardless of the specific retinol exit pathway. It could rationalize why mutations in TM6 and TM7 inhibit retinol uptake (72), as they could shift the equilibrium towards a state (JM3) in which JM is close to TM7 and cannot destabilize the IM-JML interaction to favour retinol unbinding. Furthermore, JM is adjacent in sequence to CamBP0 (66), one of the STRA6 helices that directly interact with calmodulin, suggesting a possible role of calmodulin in altering the equilibrium among JM states, and ultimately regulating retinol uptake. This observation is particularly intriguing, as the role of calmodulin still remains enigmatic, with no direct link to retinol transport being identified so far.

## Conclusions

We have reported a method to determine structure and dynamics of proteins by modelling thermodynamic ensembles from cryo-EM density maps. The application to the integral membrane receptor STRA6 illustrated how functional dynamics might remain hidden in areas of the map at lower resolution. The method is capable of revealing such dynamics by disentangling the effect of noise in the maps from conformational heterogeneity. This method is implemented in the PLUMED-ISDB module (51) of the open-source PLUMED library (www.plumed.org) (52), allowing the integration of cryo-EM with other ensemble-averaged experimental data, thus readily enabling integrative structural and dynamical biology studies (18, 19). The approach can be extended to model multiple ensembles using 3D reconstructions obtained from different 2D class-averages and to thoroughly characterize the conformational landscape, dynamics, and function of complex biological systems.

## Acknowledgement

The metainference simulations were performed at the Sisu supercomputer menaged by the CSC - IT Center for Science (Finland) under the DECI-14 project “EMMI”.

